# Engineering Conjugative Plasmids for Inducible Horizontal DNA Transfer

**DOI:** 10.1101/2025.03.12.642913

**Authors:** Tahani Jaafar, Emily Carvalhais, Arina Shrestha, Ryan R. Cochrane, Jordyn Meaney, Stephanie L. Brumwell, Samir Hamadache, Vida Nasrollahi, Bogumil J. Karas

**Author notes:** Corresponding author: Bogumil J. Karas. Co-first authors (Tahani Jaafar, Emily Carvalhais).

## Abstract

Rapidly developing microbial resistance to existing antimicrobials poses a growing threat to public health and global food security. Current chemical-based treatments target cells by inhibiting growth or metabolic function, but their effectiveness is diminishing. To address the growing antimicrobial resistance crisis, there is an urgent need for innovative therapies. Conjugative plasmids, a natural mechanism of horizontal gene transfer in bacteria, have been repurposed to deliver toxic genetic cargo to recipient cells, showing promise as next-generation antimicrobial agents. However, the ecological risks posed by unintended gene transfer require robust biocontainment strategies. In this study, we developed inducible conjugative plasmids to solve these challenges. Utilizing an arabinose-inducible promoter, we evaluated 13 plasmids with single essential gene deletions, identifying trbC and trbF as strong candidates for stringent regulation. These plasmids demonstrated inducibility in both cis and trans configurations, with induction resulting in up to a 5-log increase in conjugation efficiency compared to uninduced conditions. Although challenges such as reduced conjugation efficiency and promoter leakiness persist, this work establishes a foundation for the controlled transfer of plasmids, paving the way for safer and more effective antimicrobial technologies.

## INTRODUCTION

Antimicrobial resistance is a persistent global challenge, including the alarming capability of many organisms, such as bacteria and fungi, to develop resistance to all known antimicrobial drugs. In recent years, fungal pathogens have emerged as a significant threat due to rising resistance to antifungal treatments. Recognizing this escalating crisis, the World Health Organization (WHO) released its first Fungal Priority Pathogens List in 2022 (World Health Organization. Antimicrobial Resistance Division et al. 2022), underscoring the immense public health risks posed by fungal infections worldwide. The same challenges are evident in agriculture, where resistance amongst plant pathogenic fungi threatens global food security.

Currently, a novel class of antimicrobials is being developed based on bacterial conjugation, a natural mechanism of horizontal gene transfer requiring direct cell-cell contact (Hamilton et al. 2019; Neil et al. 2020, 2021). This process utilizes the type IV secretion system to deliver DNA, including plasmids encoding proteins that are toxic to the targeted recipient, killing the recipient cells upon delivery. Conjugation can occur in a liquid environment, on top of or within solid media (Hamilton et al. 2019; Soltysiak et al. 2019). Since bacterial conjugation can be used for delivery *in situ*, recently, there has been increased interest in repurposing it as an antimicrobial against bacterial species (López-Igual et al. 2019; Hamilton et al. 2019) as well as in mouse microbiome models (Neil et al. 2020, 2021). Cross-kingdom conjugation from bacteria to yeast and algae has also been demonstrated (Hayman and Bolen 1993; Moriguchi et al. 2013; Karas et al. 2015; Brumwell et al. 2019). Despite its potential utility, cross-kingdom conjugation remains limited due to relatively low conjugation frequency compared to bacterial recipients. However, synthetic biology approaches have shown that conjugative plasmids can be engineered to increase plasmid transfer to yeast by up to 23-fold and demonstrated engineered plasmids’ antifungal properties (Cochrane et al. 2022; Stindt and McClean 2024). As technology is being optimized to improve plasmid transfer, parallel efforts must focus on engineering biocontainment features.

Biocontainment is a critical consideration for deploying conjugative plasmids as antimicrobial agents. For these systems to be used safely and effectively, multilayered controls must be integrated to ensure precise regulation to minimize risks of exposure, misuse, or unintended release. These controls should include the following features: 1) Inducible conjugation machinery: The conjugation process should be activated only under specific, predefined conditions. Such inducible systems reduce the risk of horizontal gene transfer to unintended environments, thereby enhancing ecological safety. 2) Inducible target gene expression: Target gene expression, such as CRISPR systems or toxic genes, should be tightly regulated and linked to specific triggers. For example, metabolic regulators can be used to control expression (Pellegrino et al. 2023), or species-specific introns can be incorporated into the toxic gene to ensure proper translation only occurs in the target species (Cochrane et al. 2022). 3) Controllable conjugative plasmid elimination: Systems for controlled plasmid elimination can be engineered, such as plasmid-specific nucleases that can be triggered under defined conditions. For instance, intein-based thermoregulated meganucleases (Foo et al. 2024) provide a promising method for genetic containment. Together, these inducible features collectively serve as a robust biocontainment strategy, addressing key concerns regarding the ecological and safety implications of conjugative antimicrobial technologies.

To address one of these biocontainment needs, we explored the feasibility of creating inducible conjugative plasmids by evaluating whether conjugative plasmids with a single essential gene deletion could be complemented in trans. Using arabinose-inducible plasmids for complementation, six of the 13 plasmids complemented the deletion plasmids. However, four of these plasmids displayed complementation even without arabinose induction, likely due to promoter leakiness and low expression levels of the complementation gene. Notably, two plasmids exhibited tight regulatory control and were subjected to further detailed analysis. While we successfully demonstrated the feasibility of constructing such plasmids, the conjugation rate significantly declined, indicating the need for further optimization. This inducible system was also tested in liquid cultures and in conjugation experiments targeting yeast, demonstrating its versatile potential. These findings highlight both the promise and the challenges of engineering inducible conjugative plasmids. While key hurdles such as regulatory control and reduced conjugation efficiency remain, this work establishes a critical foundation for the safe and effective use of conjugative plasmids as next-generation antimicrobial agents.

## MATERIAL AND METHODS

### Strains and Growth Conditions

Recipient *Escherichia coli* Epi300 (Lucigen Corp., Cat #: LGN-EC300110, USA) with neomycin resistance gene chromosomally integrated were grown in Luria-Bertani (LB) media supplemented with neomycin (100 μg mL^-1^; BioBasic, Cat #: NB0366, Canada). *E. coli* containing cis plasmids were grown in LB media supplemented with gentamicin (40 μg mL^-1^; BioBasic, Cat #: GB0217, Canada). *E. coli* containing trans plasmids were grown in LB media supplemented with antibiotics selecting for both the deletion and the complementation plasmids, gentamicin (40 μg mL^-1^) and chloramphenicol (30 μg mL^-1^; BioBasic, Cat #: CB0118, Canada), respectively. All plasmids in the induced state were cultured with arabinose (100 µg mL^-1^; BioBasic, Cat #: ARB222, Canada), or with glucose (0.01 g mL^-1^; BioBasic, Cat#: GB0219, Canada) in the repressed state. Solid media for the bacterial conjugation was LB media with 1.5% agar (BioShop Canada Inc., Cat # AGA001.500, Canada) supplemented with arabinose (100 µg mL^-1^) or glucose (0.01 g mL^-1^) for the induced and repressed states respectively; the uninduced state was not supplemented. For transconjugant selection plates, the media was supplemented with gentamicin (40 μg mL^-1^) and neomycin (100 μg mL^-1^).

*Saccharomyces cerevisiae* VL6−48 (ATCC MYA-3666: MATα, his3-Δ200, trp1-Δ1, ura3-52, lys2, ade2-101, met14, psi + cir^0^) was grown in 2x yeast extract peptone dextrose media (YPD) supplemented with adenine hemisulfate (200 µg mL^-1^; Sigma-Aldrich, #A2545, St. Louis, MO, USA) (YPAD) and ampicillin (100 µg mL^-1^; BioBasic, Cat #: AB0028, Canada). For yeast conjugation experiments, the conjugation plates consisted of complete minimal (CM) glucose media lacking histidine (Teknova, Inc) with 10% LB and adenine hemisulfate (80 µg mL^-1^; Sigma-Aldrich, #A2545, St. Louis, MO, USA) in 2% agar. Selection plates consisted of CM glucose media lacking histidine supplemented with adenine hemisulfate (80 µg mL^-1^). No antibiotics were added to the conjugation nor the selection plates.

### *Trans* Plasmid Creation

Gene inserts were amplified from pTA-Mob 2.0 and the expression plasmid was amplified in three overlapping fragments using pAGE2.0i template (Supplemental Figure S1) with GXL polymerase (Takara Bio Inc., Cat #: R050A, Japan) according to the manufacturer’s instructions. Genes of interest used annealing temperatures between 57-60ºC with 25-30 cycles and backbone fragments used 60°C annealing temperature for 30 cycles. Plasmids were then assembled using four fragments via yeast-mediated cloning (primers are listed in Supplemental Table S1) and 20 colonies were passed on selection media (complete minimal glucose media lacking histidine) twice. Using Qiagen Multiplex Kit (Qiagen, Inc., Cat #: 206143, Germany), multiplex (MPX) PCR was performed on the first 10 colonies with diagnostic primers (BK888_F/R), which would result in a 214 bp amplicon if the gene insert was not incorporated. The MPX PCR conditions were as follows: a 95°C hot start for 15 min, followed by 35 cycles of 94°C for 30 s, 57°C for 90 s, and 72°C for 2 min, followed by a final extension at 72°C for 5 min, and a 12°C infinite hold. Positive MPX clones were isolated and transformed into *E. coli* by electroporation. Three clones were grown in LB supplemented with arabinose (100 µg mL^-1^) and DNA was isolated using miniprep column (BioBasic; Cat #: S614, Canada). Plasmid DNA (approximately 600 ng) was screened by diagnostic digestion using BamHI (New England Biolabs Inc, Cat #: R0136, USA) with 1 hour incubation at 37ºC. Final clones (1-2) were PCR amplified and verified by Sanger sequencing.

### *Cis* Plasmid Creation

The inducible plasmids were made based on the pSC5GGv1 plasmid created by Cochrane et al. 2022. The *traJ* sequence located downstream to the origin of transfer in the vector backbone, and the native *trbF* genes were deleted. For this, the fragment spanning *traJ* deletion was amplified using pSC5.1 as the template and the fragment with *trbF* deletion was amplified using *trbF* knockout plasmid (Cochrane et al. 2022). Inducible gene constructs were inserted at the golden-gate landing pad using methods previously described. For pSC6, the *trbF* gene along with mRFP driven by an arabinose inducible pBAD promoter and terminator were amplified out of the *trbF* inducible plasmid with primer-mediated addition of BsaI restriction sites and homology to insert downstream of the I-SceI restriction site in the pSC5 backbone. Primers and templates used to amplify all the fragments are listed in Supplemental Table S2. pSC6 was also submitted https://www.addgene.org/, ID: 233020.

### Inducible Conjugation to Bacteria – Solid Media

*Preparation of E. coli*. Donor and recipient *E. coli* were grown overnight at 37ºC with shaking at 225 rpm in 5 mL of LB media inoculated with a single colony and supplemented with the appropriate antibiotics. Saturated cultures were diluted to an optical density at 600 nm (OD_600_) of 0.1 into a 50 mL culture of LB the following day and supplemented with the appropriate antibiotics. Donor strains were grown in three states: induced, uninduced, and repressed as described above. All 50 mL cultures were grown to an OD_600_ of 1.0 and pelleted for 15 min at 3000 relative centrifugal force (RCF). After centrifugation, the supernatant was decanted and donor and recipient pellets were resuspended in 5 mL or 0.5 mL of ice-cold 10% glycerol, respectively. Aliquots with 500 μL of cells were frozen in a -80°C ethanol bath and stored at -80°C.

*Conjugation*. Induced, uninduced, and repressed conjugation plates (20 mL, 1.5% agar, LB with appropriate supplements) were dried in a biosafety cabinet for 20 minutes. Aliquots of the donor and recipient strains were thawed on ice during this time. Donor and recipient cells were added in a 1:3 ratio (33 μL:100 μL) to an Eppendorf tube and pipetted up and down before adding the total volume to the respective conjugation plate and spreading the mixture evenly. Plates were allowed to dry for 5 minutes before incubating at 37ºC for 1.5 hours without stacking. Following the conjugation period, plates were scraped with 750 μL of cold sterile ddH_2_O twice into an Eppendorf tube where the volume was adjusted to 1.5 mL and then vortexed for 5 seconds. Using a 96 well plate, ten-fold serial dilutions were performed from 10^−1^ to 10^−5^. For the conjugation in cis, 100 μL of each dilution was plated on the selection plates (20 mL, 1.5% agar, LB with gentamicin and neomycin). For the conjugation in trans, 25 μL of each dilution was mixed into 100 μL of sterile ddH_2_O and then plated onto the selection plates (20 mL, 1.5% agar, LB with gentamicin and neomycin). Plates were allowed to dry for 5 minutes before being incubated at 37ºC for 24 hours.

### Inducible Conjugation to Bacteria-Liquid Media

*Preparation of E. coli*. Donor and recipient cells were prepared according to the description above in “Inducible Conjugation to Bacteria – Solid Media”.

*Conjugation*. Frozen aliquots of the donor and recipient *E. coli* were thawed on ice for 20 minutes. In 15 mL Falcon tubes, 3 mL of LB media was prepared with or without arabinose (100 µg mL^-1^) supplementation lacking all antibiotics. Then, 500 μL of donor was added along with 50 μL of recipient to the LB media. The tubes were incubated at 37ºC with no shaking for 14 hours (Supplemental Figure S6 showcases shaking & non-shaking conditions). After the conjugation period, the liquid culture was pipetted up and down 10 times and 200 μL (undiluted) was plated onto selection plates (20 mL, 1.5% agar, LB with gentamicin and neomycin). Plates were allowed to dry for 5 minutes and before being incubated at 37ºC for 24 hours.

### Complementation Analysis

The protocol outlined in “Inducible Conjugation to Bacteria – Solid Media” section above was followed using all donor strains created harbouring trans plasmids. However, 75 μL of each dilution were plated on the selection plates which were then incubated at 37ºC for 48 hours.

### Inducible Conjugation to Yeast

*Preparation of E. coli and S. cerevisiae*. Donor *E. coli* were grown overnight at 37ºC with shaking at 225 rpm in 5 mL of LB media inoculated with a single colony and supplemented with the appropriate antibiotic. Recipient *S. cerevisiae* was grown at 30ºC with shaking at 225 rpm in 5 mL of 2x YPAD supplemented with ampicillin (100 µg mL^-1^). Saturated cultures were diluted to an OD_600_ of 0.1 into a 50 mL culture of LB or 2x YPAD the following day and supplemented with the appropriate antibiotics. *E. coli* strains were grown in three states: induced, uninduced, and repressed as described above. Donor and recipient cultures were grown to an OD_600_ of 1.0 or 3.0, respectively, and pelleted for 15 min at 3000 RCF. After centrifugation, the supernatant was decanted and donor and recipient pellets were resuspended in 0.5 mL and 1 mL of ice-cold 10% glycerol, respectively. Then, 500 μL aliquots were frozen in an ethanol bath and stored at -80°C.

*Conjugation*. Induced, uninduced, and repressed conjugation plates (20 mL, 1.5% agar, 10% LB medium, CM glucose broth lacking histidine with appropriate supplement) were dried in a biosafety cabinet for 20 minutes. Aliquots of the donor and recipient were thawed on ice during this time. Then, 100 μL of both donor and recipient cells were added into an Eppendorf tube and pipetted up and down before adding the total volume to the respective conjugation plate and spreading the mixture evenly. Plates were allowed to dry for 5 minutes before incubating at 30ºC for 3 hours without stacking. Following the conjugation period, plates were scraped with 2 mL of cold sterile ddH_2_O and readjusted in a 15 mL Falcon tube and then vortexed for 5 seconds. Two technical replicates were performed with 200 μL and 400 µL of the undiluted cells spread on selection plates (30 mL, 1.5% agar, CM glucose broth lacking histidine). Plates were allowed to dry for 5 minutes before being incubated at 30ºC for 5 days.

### Statistical Analysis

The pairwise comparisons between groups were made using Student’s *t*-test with either equal or unequal variance based on the result of an *F*-test. Data were expressed as mean ± standard error of the mean of at least four biological replicates. The tests were considered statistically significant when *P* <0.05 (*).

## RESULTS

To make the *in trans* inducible conjugative plasmid, we used a previously generated resources, a single gene deletion library of conjugative plasmid pTA-Mob 2.0 (Cochrane et al. 2022) and constructed corresponding complement plasmids. Complement plasmids contained a single gene driven by the pBAD promoter. This promoter is arabinose-inducible; transcription of genes driven by the pBAD promoter can be induced by the addition of arabinose and repressed by the addition of glucose. Based on our published results (Cochrane et al. 2022), we selected 13 plasmids with single gene deletions essential for conjugation from *E. coli* to *S. cerevisiae* (**Table 1**). We transferred these plasmids and their corresponding complement plasmids to an *E. coli* donor and performed conjugation experiments on solid media (**Table 1**, Supplemental Figure S2). Since both plasmids (pTA-Mob 2.0 deletions and complement plasmids) have origins of transfer, if complementation occurs, one and/or the other can be transferred to recipient cells, as depicted schematically in **Figure 1**, Supplemental Figure S3.

**Table 1.** Conjugation complementation results for 13 plasmids, each with a single essential gene deletion. “Yes” indicates successful conjugation (transconjugants obtained), demonstrating that the deleted gene was effectively complemented. “No” indicates failed conjugation (no transconjugants obtained); The red and green colors indicate undesired and desired outcomes, respectively. Specifically, the desired outcome for induced condition is Yes and for the repressed and uninduced conditions is No. See Supplementary Figure S2.

| Deleted Gene | Induced | Repressed | Uninduced |
| --- | --- | --- | --- |
| *trbA* | No | Not Tested | Not Tested |
| *trbC* | Yes | No | No |
| *trbD* | Yes | Yes | Not Tested |
| *trbF* | Yes | No | No |
| *trbI* | No | Not Tested | Not Tested |
| *trbJ* | No | Not Tested | Not Tested |
| *trbL* | Yes | Yes | Not Tested |
| *traF* | Yes | Yes | Not Tested |
| *traG* | Yes | Yes | Not Tested |
| *traH* | No | Not Tested | Not Tested |
| *traJ* | No | Not Tested | Not Tested |
| *traK* | No | Not Tested | Not Tested |
| *traM* | No | Not Tested | Not Tested |

**Figure 1.**
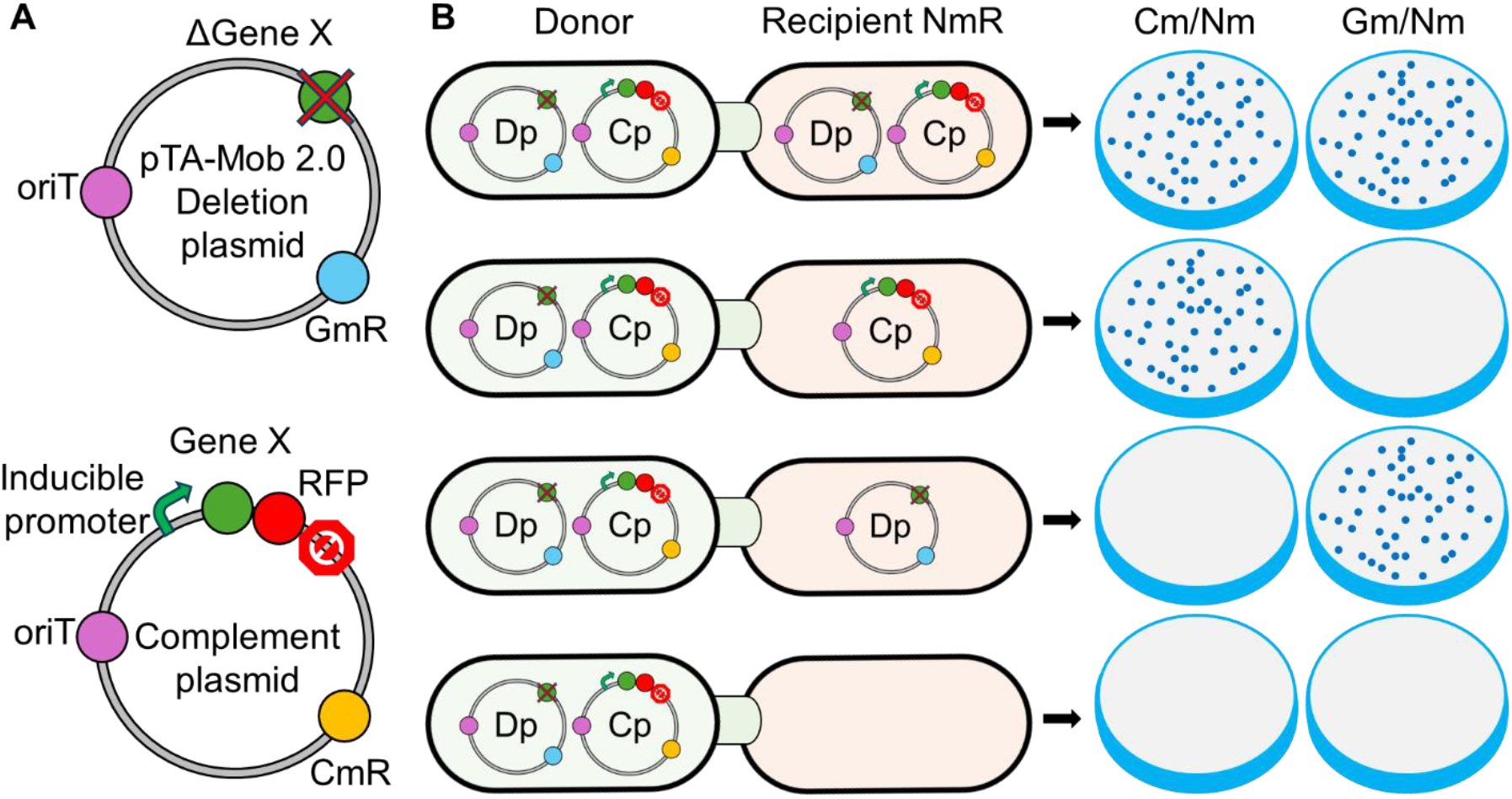
Schematic of plasmids used in trans-conjugation and possible outcomes. A) Single-gene deletion plasmids (Dp) (Cochrane et al. 2022) were derived from the conjugative plasmid pTA-Mob 2.0 (Soltysiak et al. 2019). A key feature of the Dp is the presence of all necessary conjugative machinery for plasmid mobilization and transfer, except for one essential conjugation gene. The plasmid also contains a selectable marker for *E. coli*, providing gentamicin resistance (GmR; Blue). The complement plasmid (Cp) includes the essential conjugation gene (Gene X; Green) that was removed from the Dp under the control of an inducible pBAD promoter. Directly downstream of this gene is a red fluorescent protein (RFP; ed) reporter. The selection marker for Cp in *E. coli* provides chloramphenicol resistance (CmR; Orange). B) Schematic of possible outcomes following *E. coli* to *E. coli* conjugation. Recipient cells have a chromosomally integrated neomycin resistance marker (NmR). Using double-antibiotic selection plates with chloramphenicol and neomycin or gentamicin and neomycin, it is possible to evaluate whether one, both, or neither plasmid was transferred.

Our initial experiments showed that conjugation was restored for 6 out of 13 deletion plasmids, but only two, *trbC* and *trbF*, showed full repression of conjugation when glucose was added. Even without repression (uninduced condition), we did not observe conjugation for these genes (**Table 1**). Therefore, we have identified two strong gene candidates to form the basis of our next-generation inducible conjugation plasmids.

To further evaluate this *in trans* conjugation setup, we conducted a larger experiment involving four biological replicates for the *trbF* and *trbC* deletion/complement plasmid combinations to an E. coli recipient. Our findings confirmed the initial results, demonstrating that the conjugative system is indeed inducible (**Figure 2, Supplemental Figure S4, Supplemental Table S3**). However, conjugation efficiency was notably lower compared to the parental plasmid pTA-Mob 2.0, showing a decrease of almost two orders of magnitude. While the system exhibited some leakiness, induction led to a significant enhancement in conjugation efficiency, producing approximately four orders of magnitude more transconjugant colonies as compared to uninduced conditions.

**Figure 2.**
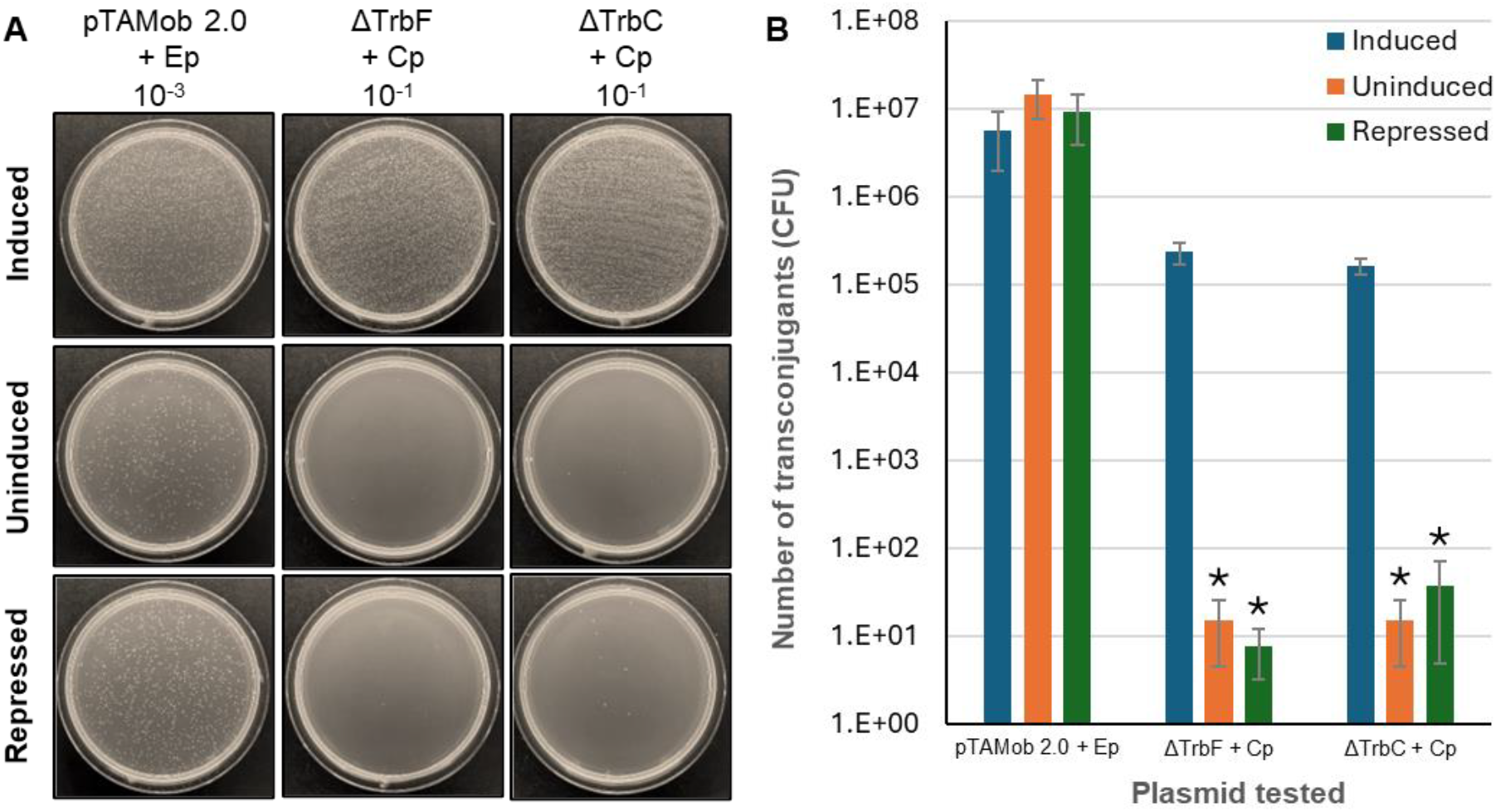
Analysis of the inducible trans-conjugative system. A) Representative transconjugant colonies on final selection plates (Nm/Gm). The donor strains included pTA-Mob 2.0 with an empty plasmid (Ep) and pTA-Mob 2.0 carrying deletions of the *trbF* (ΔTrbF) or *trbC* (ΔTrbC) genes, complemented with their respective Cp plasmids. All combinations were tested under three conditions: induced with arabinose, uninduced, and repressed with glucose. Complete figures of all replicates displayed in Supplemental Figures S4. B) Average colony counts for all three conditions across all replicates. Complete colony counts for all replicates displayed in Supplemental Table S3. Error bars represent the standard error of the mean (SEM). Student’s t-test: ^*^p < 0.05; n = 4.

Building on the findings with inducible conjugative plasmids in trans, we designed a novel plasmid, pSC6, based on pSC5GGv1 (Cochrane et al. 2022). In this construct, the *trbF* gene was deleted and reintroduced under an inducible promoter and terminator to minimize unintended activation of adjacent genes or operons (**Figure 3**). As with the *trans* system, our *cis* conjugative system demonstrated clear inducibility. However, conjugation efficiency was significantly reduced compared to the control plasmid pSC5GGv1, exhibiting an approximately two-order-of-magnitude decrease (see **Figure 3**). Notably, induction substantially boosted conjugation efficiency, achieving a three-order-of-magnitude increase relative to uninduced conditions (**Figure 3**).

**Figure 3.**
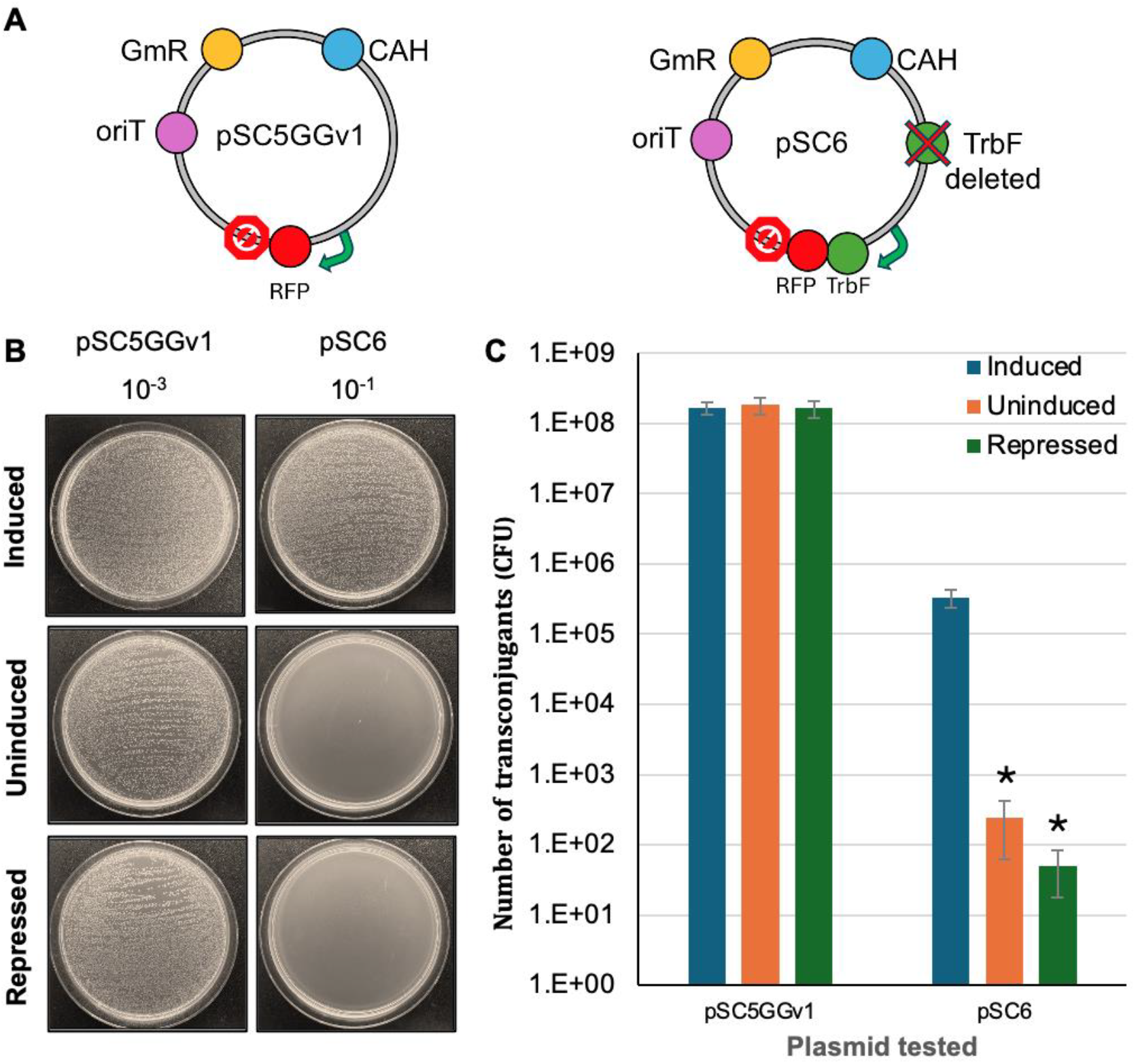
Analysis of the inducible *cis* conjugative system. A) Schematic of plasmids used in cis-conjugation. The conjugative plasmid pSC5GGv1 served as a positive control. Plasmid pSC6 had the *trbF* gene deleted and relocated to a different position under the pBAD inducible promoter. All plasmids contain a gentamicin selectable marker (GmR; yellow). CAH – elements for replication and maintenance in yeast, including centromere (CEN6), autonomously replication sequence (ARS4), and histidine auxotrophic marker (HIS3). B) Representative final selection plates (Nm/Gm) transconjugant colony frequency. The donor strains had only one plasmid each: pSC5GGv1 or pSC6. Recipient cells have a chromosomally integrated neomycin resistance marker (NmR). All plasmids were tested under three conditions: induced with arabinose, uninduced, and repressed with glucose. Complete figures of all replicates displayed in Supplemental Figure S5. C) Average colony counts for all three conditions across all replicates. Complete colony counts for all replicates displayed in Supplemental Table S4. Error bars represent the standard error of the mean (SEM). Student’s t-test: ^*^p < 0.05; n = 6.

Finally, we conducted preliminary experiments to evaluate conjugation in liquid culture from *E. coli* to *E. coli*. Our results demonstrated that pSC6 was inducible in this environment, but the conjugation efficiency was significantly lower compared to the control plasmid (see **Figure 4**). Given our goal of utilizing this inducible plasmid in antifungal technology, we further tested its ability to conjugate into yeast. pSC6 resulted in only eight colonies under induced conditions, with no colonies observed without induction. While these findings highlight the potential for inducible control of the conjugative system, further engineering will be necessary to achieve conjugation efficiencies comparable to the control plasmid.

**Figure 4.**
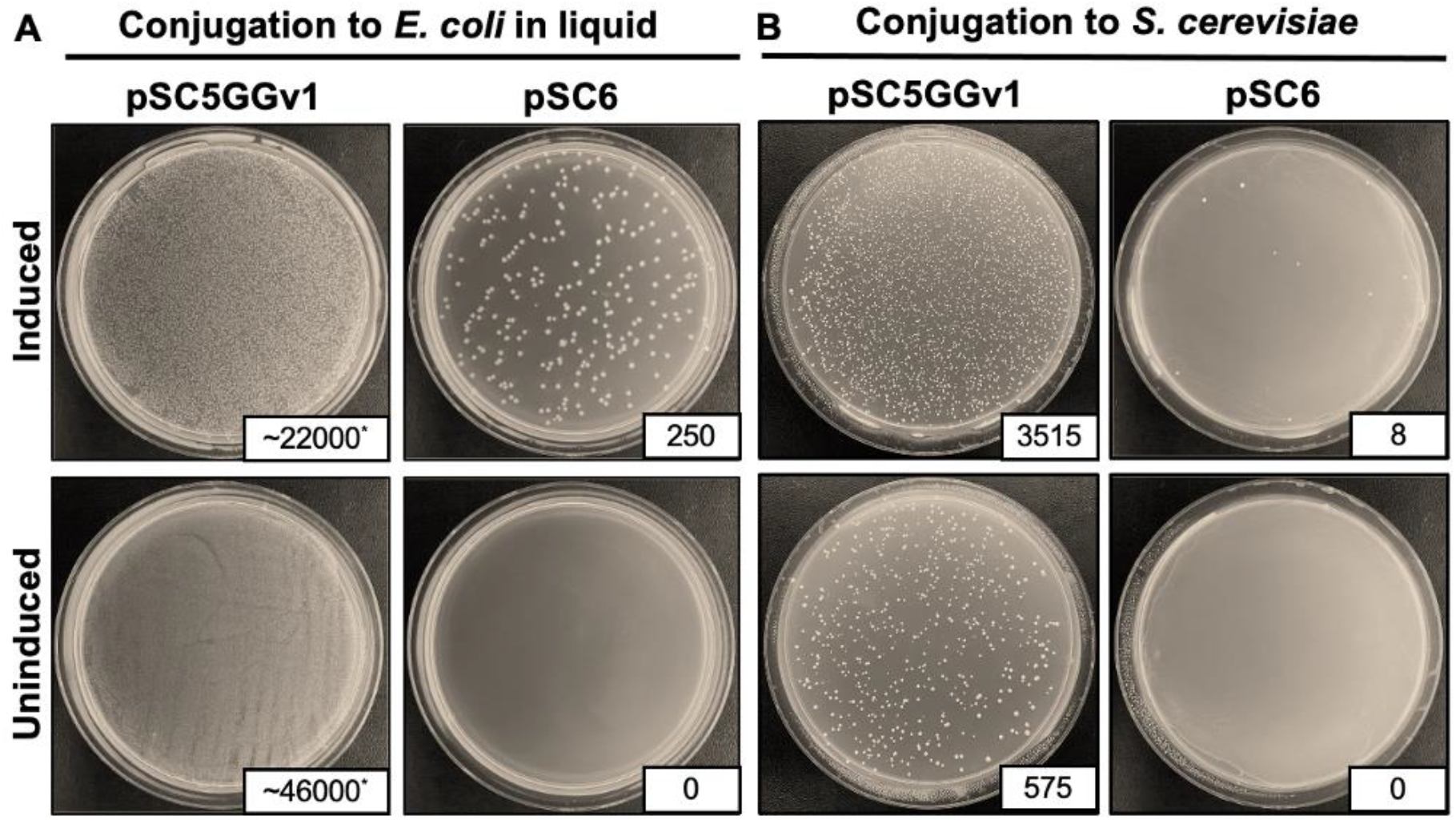
Applications of inducible conjugative plasmid. A) Representative transconjugant colonies on final selection plates (Nm/Gm) following cis-conjugation from *E. coli* to *E. coli* in liquid media. Recipient cells have a chromosomally integrated neomycin resistance marker (NmR). B) Representative transconjugant colonies on final selection plates (−HIS) following cis-conjugation from *E. coli* to *S. cerevisiae*. All plasmids include elements for selection and replication in yeast (HIS3 gene for complementation of yeast auxotrophy). ^*^colony counts were estimated by counting within a 0.5 × 0.5 cm square, and adjusted to plate size assuming equal distribution.

## DISCUSSION

In recent years, the development of conjugation-based antimicrobials has attracted significant interest. However, as this technology advances, its limitations, particularly one concerning the control of engineered plasmids once released into the environment, need to be addressed. Creating inducible conjugative systems is a fundamental requirement in horizontal gene transfer technologies. A significant example of such systems is the work by Brophy et al., who engineered the ICEBs1 system from *Bacillus subtilis* (Brophy et al. 2018). This integrative and conjugative element (ICE) facilitates efficient and inducible DNA transfer to undomesticated bacteria. Their approach involved separating the type IV secretion system from the transferred DNA, minimizing unintended genetic dissemination.

In our study, we aimed to develop a *cis* conjugative plasmid encompassing all components necessary for conjugation while maintaining stringent control. We examined 13 essential genes required for the conjugation of pTA-Mob 2.0 plasmid, including those involved in pilus formation, assembly, and stability (trbC, trbD, trbF, trbI, trbJ, trbL, traF), components of the relaxosome complex (traH, traJ, traK, traM), and the coupling of the relaxosome to the mating pair formation system (traG). Deletions of genes associated with pilus formation and stability were generally amenable to complementation *in trans*, whereas those in the relaxosome complex were not. This discrepancy may stem from the deletions disrupting regulatory elements of adjacent genes. Future efforts could focus on creating point mutations instead of full deletions to preserve regulatory control. We identified two candidates, trbC and trbF, that exhibited effective complementation when expressed using the pBAD promoter. It is likely that trbC and trbF require significantly higher expression levels to fully restore their respective conjugative functions. Consequently, the low basal expression caused by promoter leakiness may be insufficient to rescue the phenotype, making these genes particularly well-suited for stringent inducible control. However, we predict that with precise tuning of gene expression, all complemented genes (including trbD, trbL, traF, traG) could be harnessed for this approach.

Our findings allowed us to evaluate both *cis* and *trans* systems. In both configurations, we achieved inducibility, albeit with a decrease in conjugation rates. This reduction might result from altered expression of genes surrounding *trbC* and *trbF*. As discussed above, addressing this issue could involve introducing partial gene deletions or point mutations and assessing the impact of varying expression levels of the candidate genes on restoring conjugation efficiency.

Incorporating inducible promoters, such as the arabinose-inducible pBAD promoter, adds a necessary layer of biocontainment. This design ensures that conjugation occurs only under specific conditions, mitigating ecological risks associated with unintended horizontal gene transfer. Identifying *trbC* and *trbF* as tightly regulated components provides a good foundation for future inducible plasmid designs. However, challenges like promoter leakiness and reduced conjugation efficiency will need to be addressed for practical applications. Furthermore, environment and species-specific inducible promoters need to be tested. The capability to deliver plasmids to yeast highlights the potential of these systems for antifungal applications. However, the observed low conjugation efficiency in cross-kingdom transfer will need to be addressed, including the approaches described above. Additionally, laboratory accelerated evolution (Neil et al. 2021) can be performed to increase conjugation frequency. Overall, this study establishes a foundational framework for inducible conjugative plasmid technologies while highlighting challenges and opportunities for innovation in antimicrobial and genetic engineering applications.

## AUTHOR CONTRIBUTIONS

TJ: formal analysis, investigation, methodology, writing – original draft, writing – review & editing; EC: formal analysis, investigation, methodology; AS: formal analysis, investigation, methodology, writing – review & editing; RRC: formal analysis, investigation, methodology, writing – review & editing; JM: investigation; writing – review & editing; SB: investigation; writing – review & editing; SH: investigation; writing – review & editing; VN: investigation; writing – review & editing; BJK: conceptualization, formal analysis, funding acquisition, methodology, resources, supervision, writing – original draft, writing – review & editing.

## DATA AVAILABILITY

Data generated or analyzed during this study are available in the published article and its supplementary materials.

