## Supplemental file for "Engineering Conjugative Plasmids for Inducible Horizontal DNA Transfer"

*Co-first authors

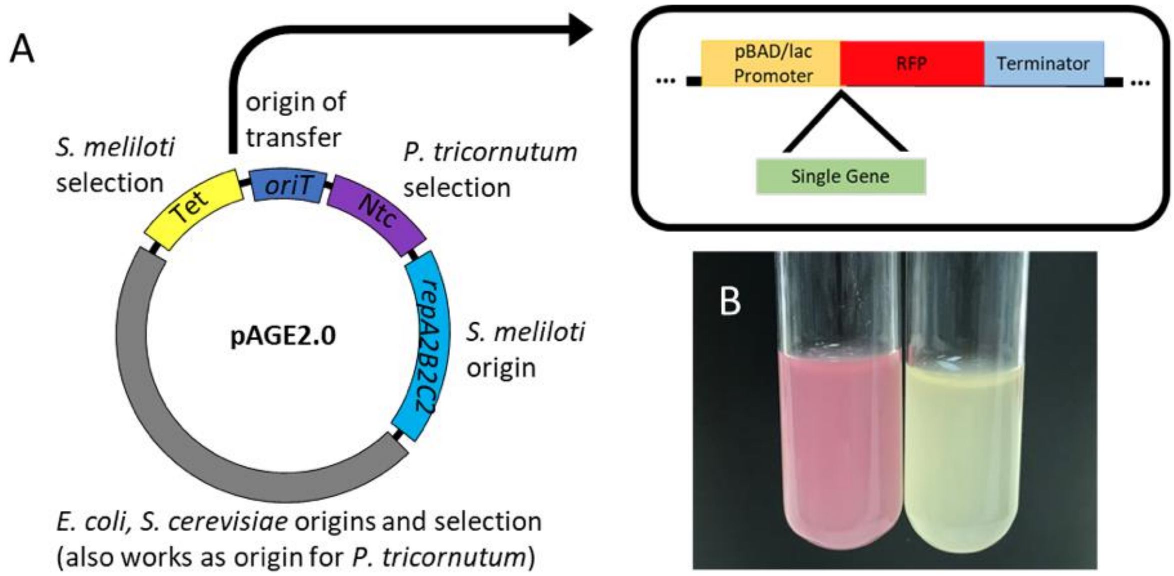


**Figure S1**. Design of inducible plasmid system – pAGE2.0i. A) Design of an inducible plasmid system using the pAGE2.0 (Brumwell et al., 2019) backbone and a pBAD or lac promoter driving the expression of RFP. Following confirmation, single genes from pTA-Mob 2.0 were added to this cassette to test overexpression following plasmid induction. B) An *E. coli* clone harbouring pAGE2.0i grown with induction by arabinose following assembly in yeast (left) and without induction (right). Note – this plasmid was mentioned in our previous manuscript (Cochrane et al., 2022) but how it was made was never described.

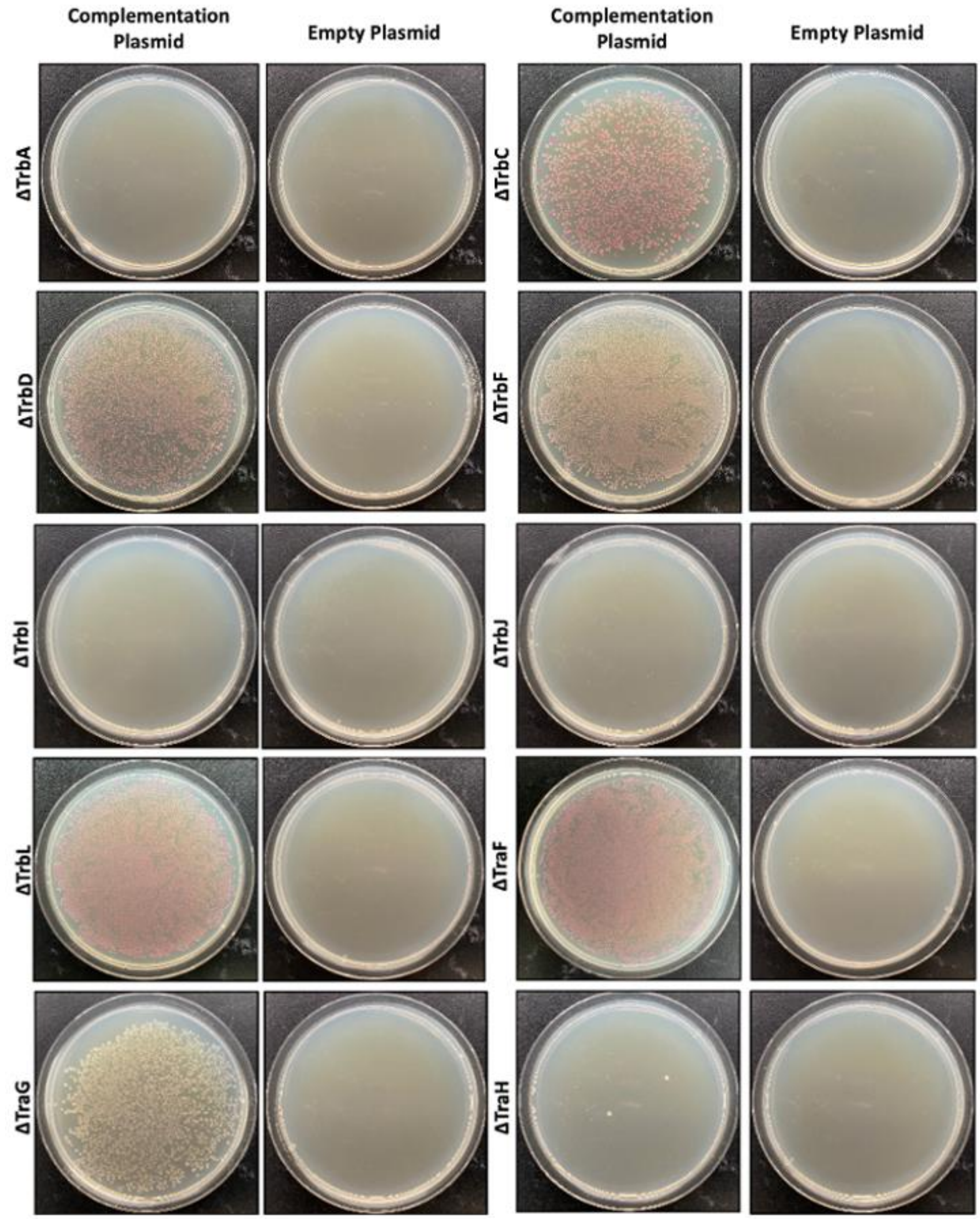
Complementation Plasmid
Empty Plasmid
Complementation Plasmid
Empty Plasmid
ΔTrbA
ΔTrbC
ΔTrbD
ΔTrbF
ΔTrbI
ΔTrbJ
ΔTrbL
ΔTraF
ΔTraG
ΔTraH

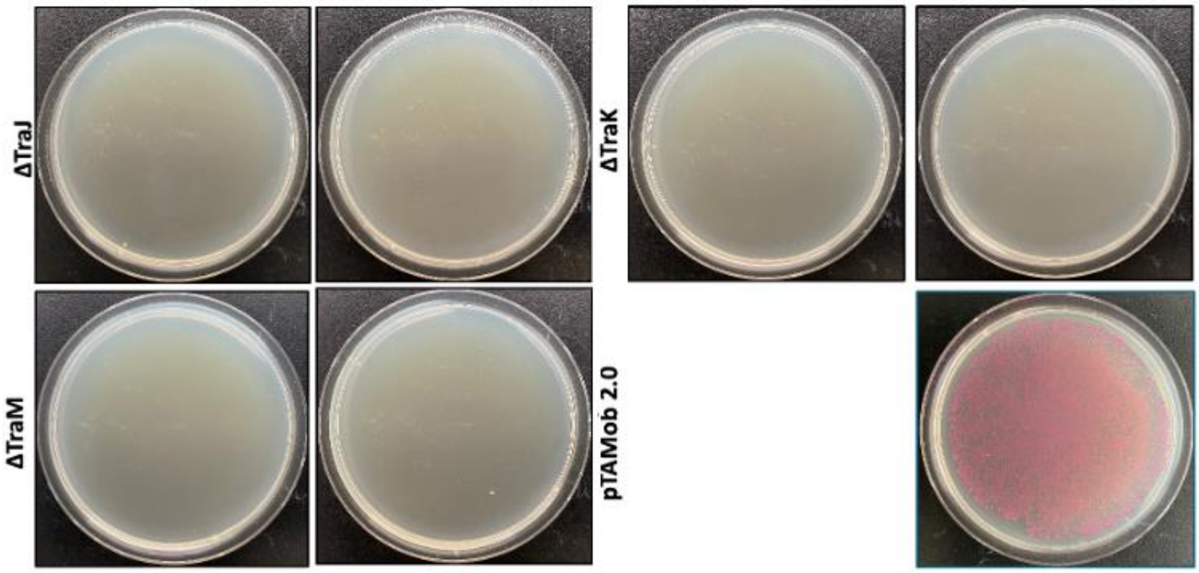


**Figure S2.** Initial complementation results for the 13 deletion plasmids listed in Table 1. Complementation and empty plasmids were tested for each gene. Complementation is successful when there is both growth on deletion plasmid + CP and no growth on deletion plasmid + EP. pTAMob 2.0 + EP was used as a control.

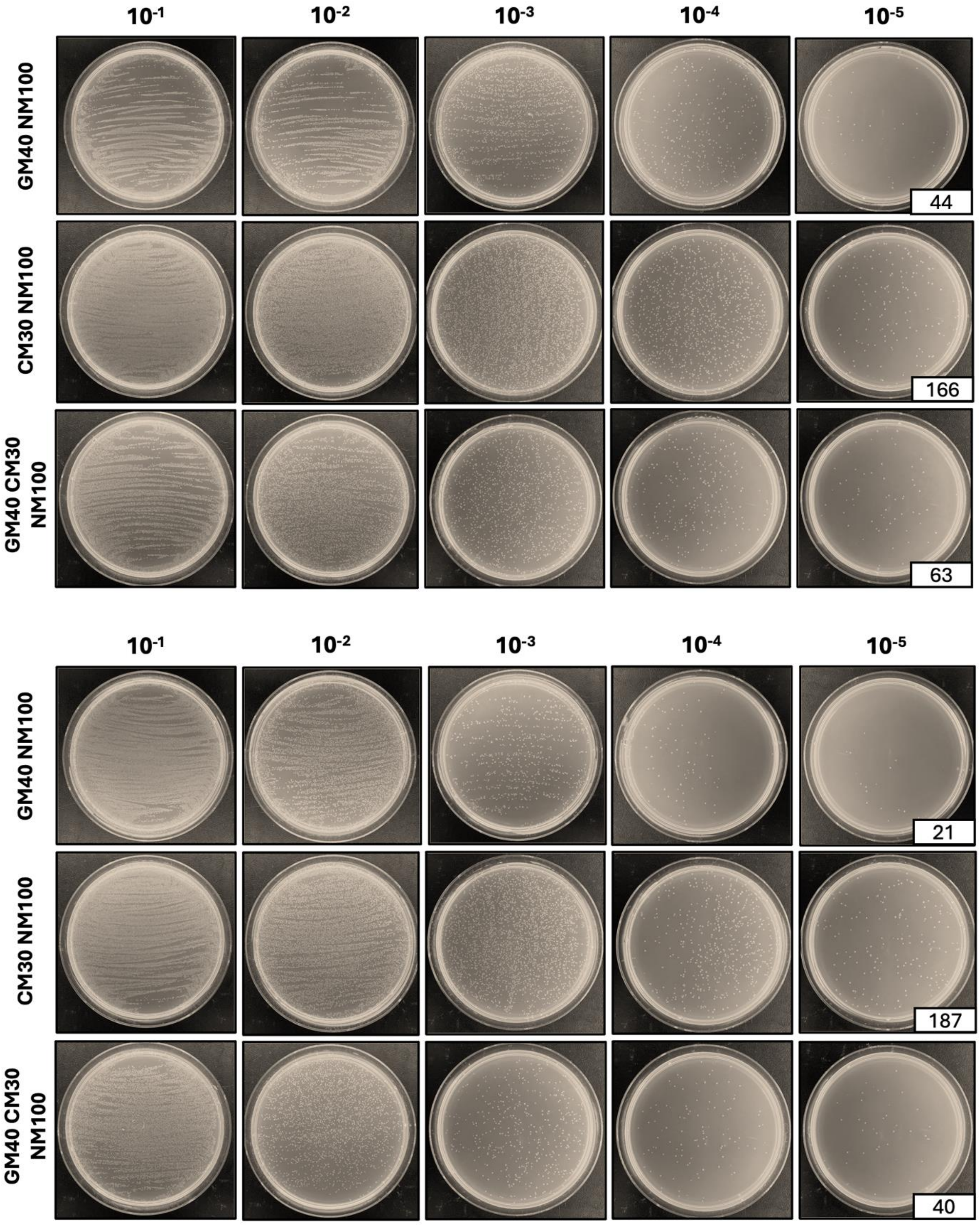


**Figure S3.** Two replicates of pTAMob 2.0 + EP conjugation in uninduced state plated on 3 different selection plates to present all possible outcomes. Selection on GM40NM100 shows transfer of the deletion plasmid. Selection on CM30NM100 shows transfer of the complementation plasmid. Selection on GM40CM30NM100 shows transfer of both plasmids. Colony counts presented for the $10^{-5}$ dilution.

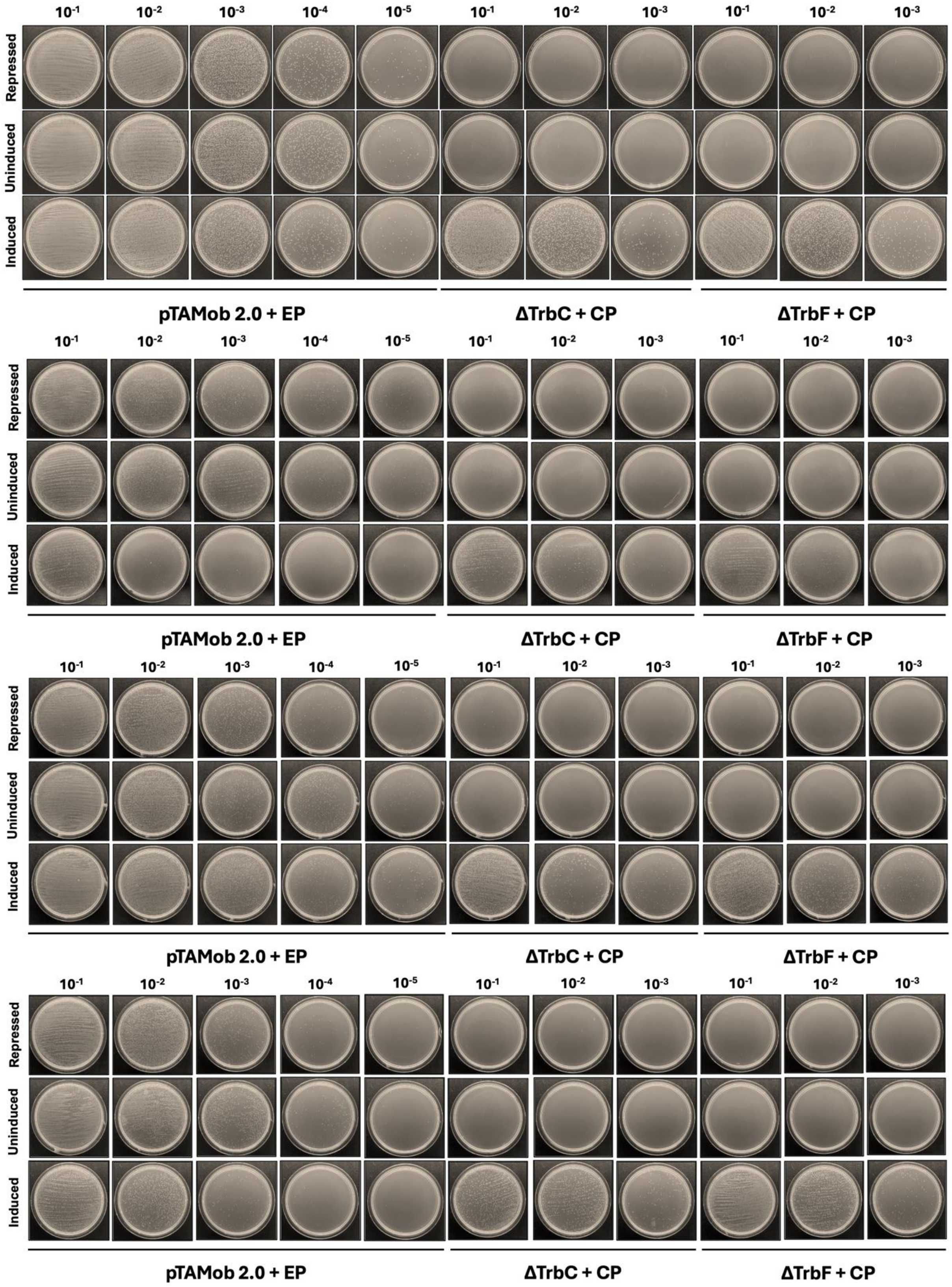


**Figure S4.** Four replicates of the inducible conjugative system in *trans*. Dilutions plated up to $10^{-5}$ to count single colonies. Colony counts presented in Table S3.

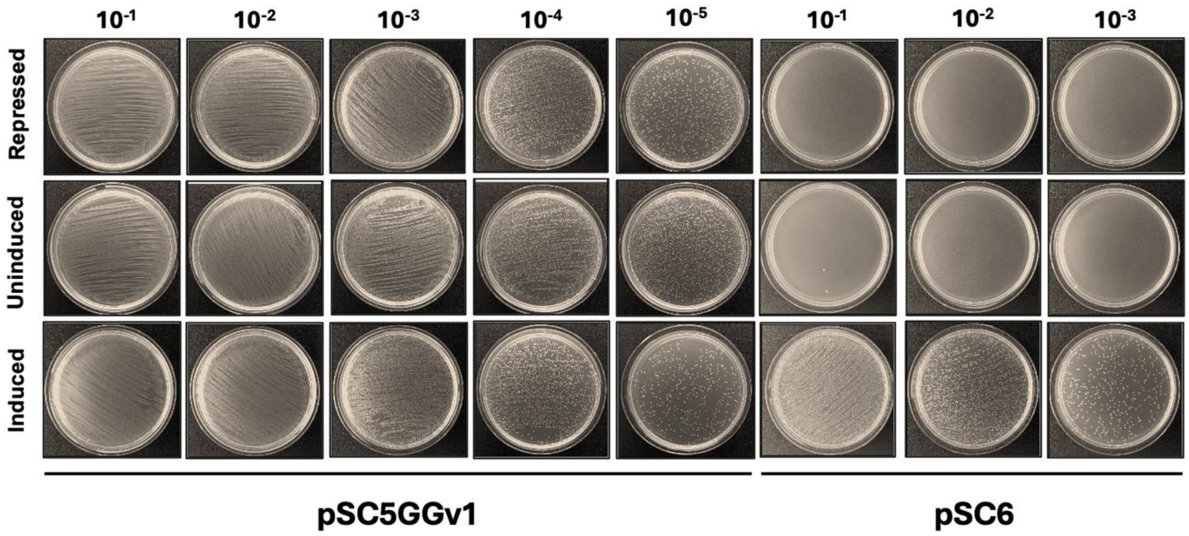
10-1
10-2
10-3
10-4
10-5
10-1
10-2
10-3
Repressed
Uninduced
Induced
pSC5GGv1
pSC6

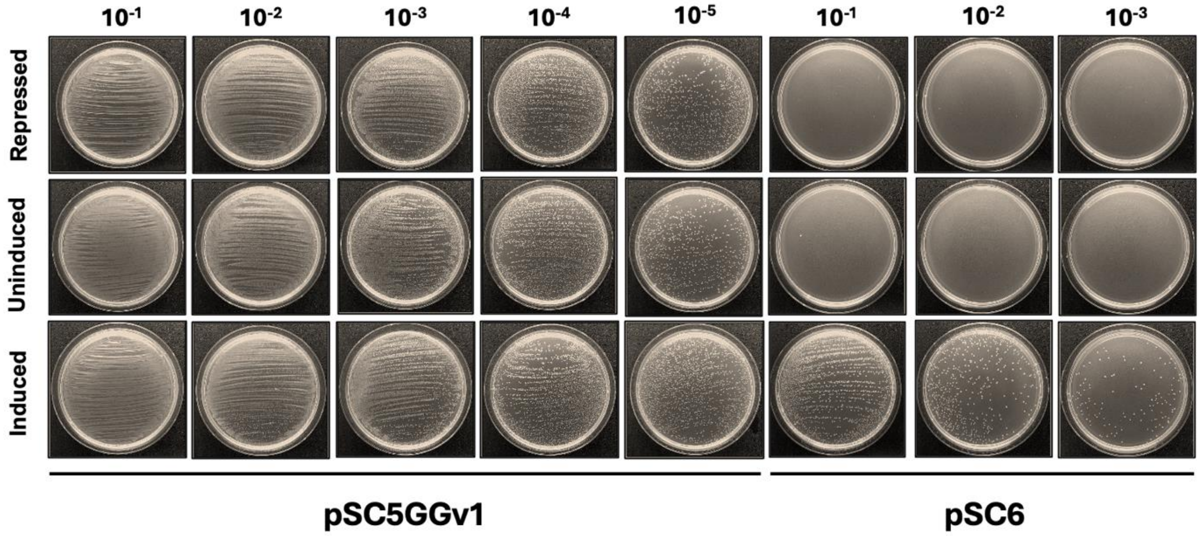
10-1
10-2
10-3
10-4
10-5
10-1
10-2
10-3
Repressed
Uninduced
Induced
pSC5GGv1
pSC6

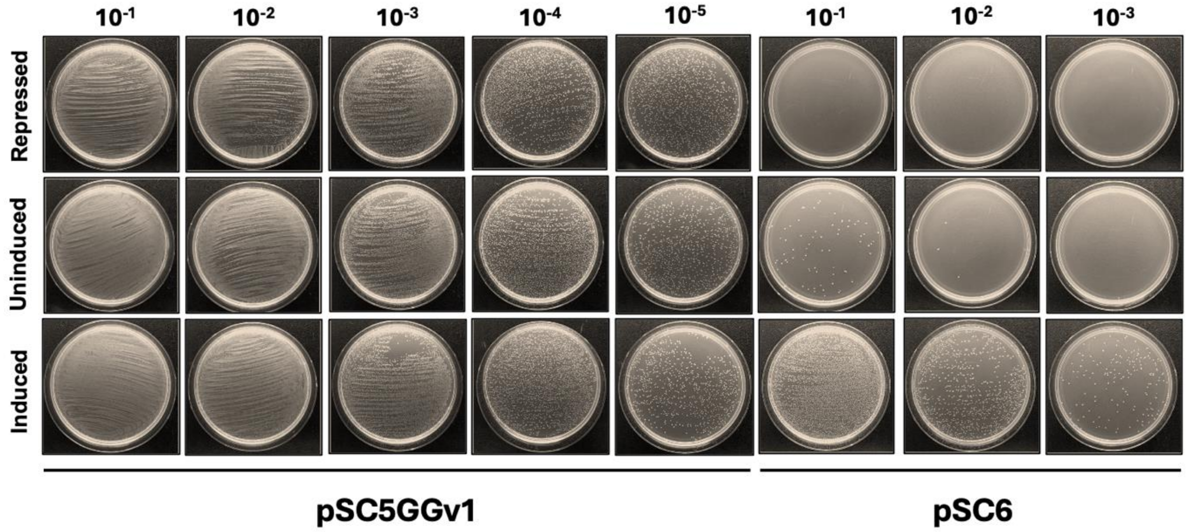
10-1
10-2
10-3
10-4
10-5
10-1
10-2
10-3
Repressed
Uninduced
Induced
pSC5GGv1
pSC6

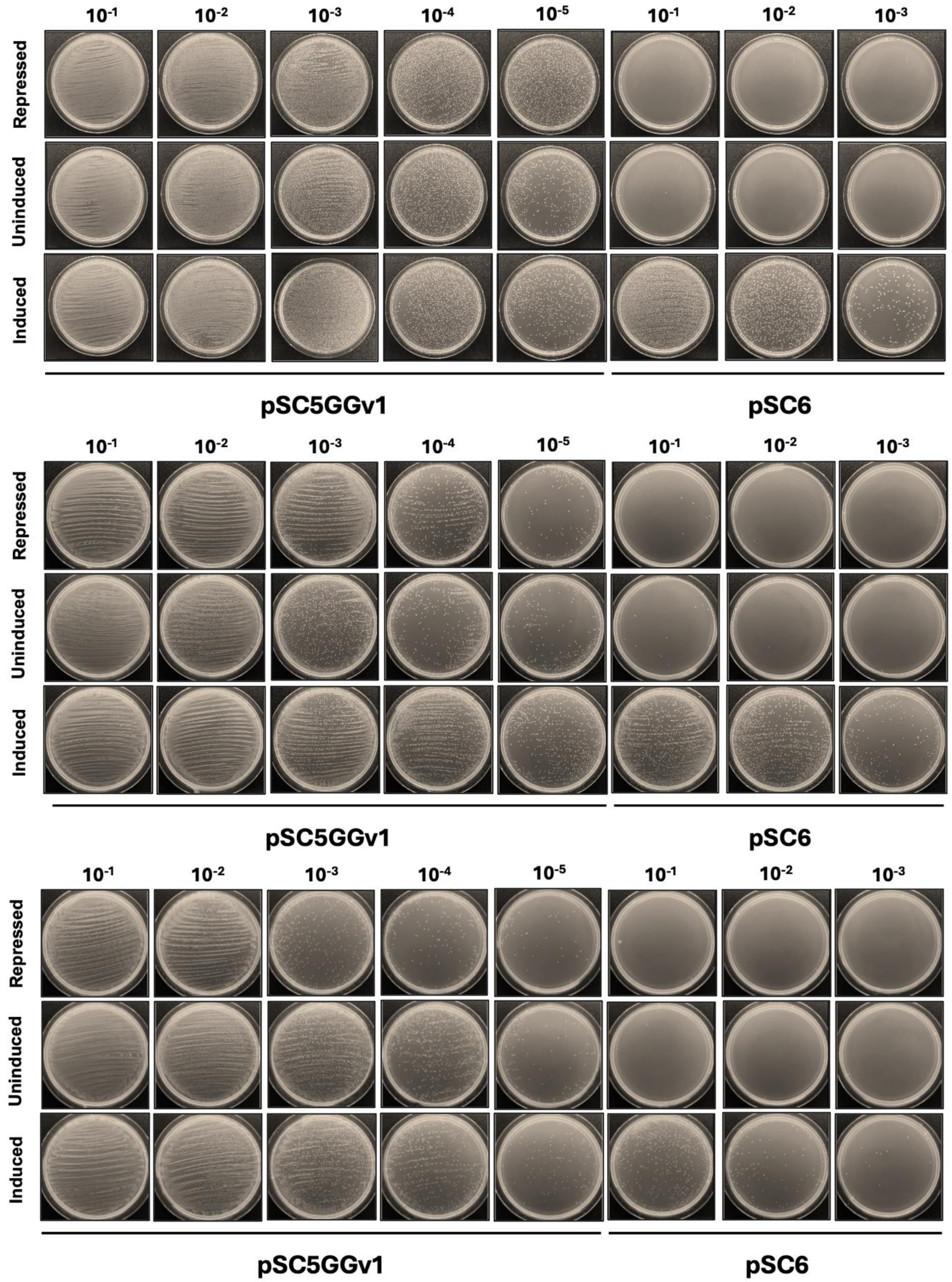


**Figure S5.** Six replicates of the inducible conjugative system in *cis*. Dilutions plated up to $10^{-5}$ to count single colonies.

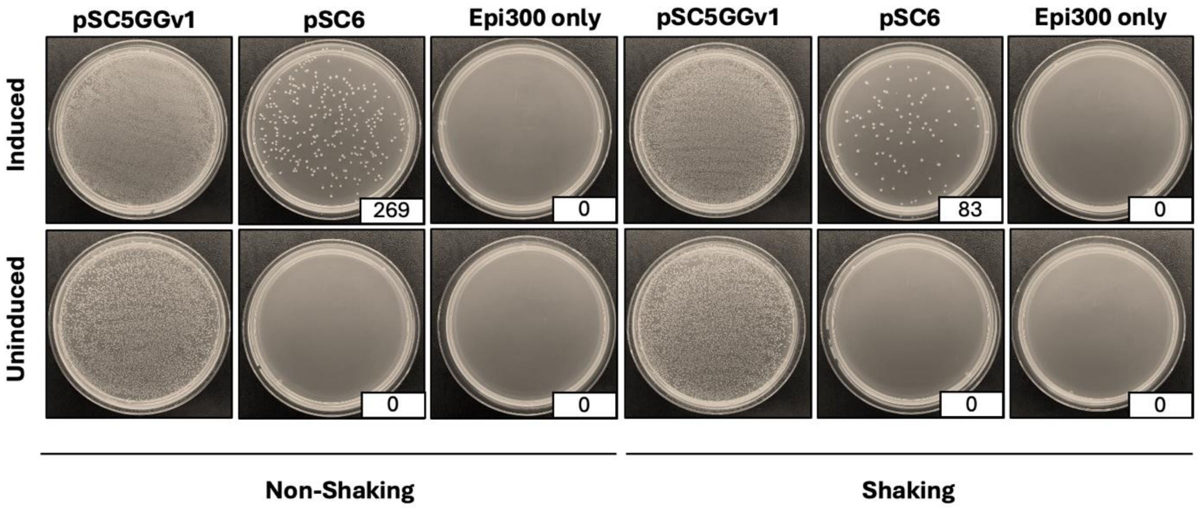


**Figure S6.** Testing conjugation in liquid in non-shaking and shaking conditions. 100uL of donor and 10uL of recipient added to 3mL of LB supplemented with arabinose in the induced state. Cultures were grown at 37ºC with or without shaking (225 rpm) for 14 hours. 200uL of culture was plated on selection plates (GM40NM100).

**Table S1.** Primers used to amplify gene inserts from pTA-Mob 2.0 and pTA-Mob 2.0i.

| Gene | Forward Primer | Reverse Primer |
|---|---|---|
| *trbA* | tttatcgcaactctctactgtttctccatacccgttttttgtttaactttaagaaggagaatgacgaaacatgagctgtc | agaaccccgcatatgtatatctccttcttaaagttaaacaaaattatttctagcccaaaatcagagccttccacgcagct |
| *trbC* | tttatcgcaactctctactgtttctccatacccgttttttgtttaactttaagaaggagaatgacaacggctgttccgtt | agaaccccgcatatgtatatctccttcttaaagttaaacaaaattatttctagcccaaaattaggcgagccgtccagccg |
| *trbD* | tttatcgcaactctctactgtttctccatacccgttttttgtttaactttaagaaggagaatggctctgcgcacgatccc | agaaccccgcatatgtatatctccttcttaaagttaaacaaaattatttctagcccaaaatcatcggtattgcttccctt |
| *trbF* | tttatcgcaactctctactgtttctccatacccgttttttgtttaactttaagaaggagaatgagttttgcagacacgat | agaaccccgcatatgtatatctccttcttaaagttaaacaaaattatttctagcccaaaatcacagaagtctcgaccagg |
| *trbI* | tttatcgcaactctctactgtttctccatacccgttttttgtttaactttaagaaggagaatgagcgaagatcaaatggc | agaaccccgcatatgtatatctccttcttaaagttaaacaaaattatttctagcccaaaattaatagtcaaacgcctggt |
| *trbJ* | tttatcgcaactctctactgtttctccatacccgttttttgtttaactttaagaaggagaatgaagaagctcgctaagaa | agaaccccgcatatgtatatctccttcttaaagttaaacaaaattatttctagcccaaaatcaccaggtcttagacgggc |
| *trbL* | tttatcgcaactctctactgtttctccatacccgttttttgtttaactttaagaaggagaatgaaaatccagactagagc | agaaccccgcatatgtatatctccttcttaaagttaaacaaaattatttctagcccaaaatcaggattgggcgggcttgt |
| *traF* | tttatcgcaactctctactgtttctccatacccgttttttgtttaactttaagaaggagaatgagccgcttccagcgcct | agaaccccgcatatgtatatctccttcttaaagttaaacaaaattatttctagcccaaaatcaccaggtcagaaccggcc |
| *traG* | tttatcgcaactctctactgtttctccatacccgttttttgtttaactttaagaaggagaatgaagaaccgaaacaacgc | agaaccccgcatatgtatatctccttcttaaagttaaacaaaattatttctagcccaaaatcatatcgtgatcccctccc |
| *traH* | tttatcgcaactctctactgtttctccatacccgttttttgtttaactttaagaaggagaatgagcaacccgaacgaaat | agaaccccgcatatgtatatctccttcttaaagttaaacaaaattatttctagcccaaaattagccctcttcctggccca |
| *traJ* | tttatcgcaactctctactgtttctccatacccgttttttgtttaactttaagaaggagaatggctgatgaaaccaagcc | agaaccccgcatatgtatatctccttcttaaagttaaacaaaattatttctagcccaaaatcatggctctgccctcgggc |
| *traK* | tttatcgcaactctctactgtttctccatacccgttttttgtttaactttaagaaggagaatgccaaagagctacaccga | agaaccccgcatatgtatatctccttcttaaagttaaacaaaattatttctagcccaaaattacagtagatcctttttgt |
| *traM* | tttatcgcaactctctactgtttctccatacccgttttttgtttaactttaagaaggagaatgagcgaccagattgaaga | agaaccccgcatatgtatatctccttcttaaagttaaacaaaattatttctagcccaaaatcataacgaggcccacacca |

**Table S2.** Primers for pSC6 creation.

| Fragment | Forward Primer | Reverse Primer | Amplicon Size (bp) |
|---|---|---|---|
| 1 | tgccgccgcgcgcatggtcgtaatgggaccgatagcccgt | tttaacctacttcctttggttccggggggatctcgcgactc | 6757 |
| 2 | atcgaagagaagcaggacga | tgctggtccatgaagatgaa | 6177 |
| 3 | tcgagctgatgtttgacgac | ggacttgaggttgctctgct | 5382 |
| 4 | ggagggaggatacaggcgaa | atcggcgtgaagcccaacagggcca | 6000 |
| 5 | gtggacattggtttcagcaa | caatgtctgatgcaatatggacaattggtttcttggtctcattaccctgttatccctaca | 6265 |
| 6 | accttcgggtgggcctttctgcgtttataggtctcatgcttacgctctagttctccttag | aagcgatgaatgatcccaag | 3477 |
| 7 | tgtaacgcttcccggtagtc | aggcccttgccaatgaat | 12097 |
| 8 | ttctttgaatgcgcgggcgtcctggtgagcgtagtccagc | agccctcccgtatcgtagtt | 10554 |
| 9 | attgaaagggtgtttgtagggataacagggtaatgagaccaagaaaccaattgtccatat | gagaacctactaaggagaactagagcgtaagcatgagacctataaacgcagaaaggccca | 1978 |
| 10 | gatgcccagcctcggtcatatttttcatggtctcatgcttacgctctagttctccttag | aagcgatgaatgatcccaag | 3477 |
| 11 | cacactggctcaccttcgggtgggcctttctgcgtttatattacagcggggtgctgcccg | gagaacctactaaggagaactagagcgtaagcatgagaccatgaaaaaatatgaccgagg | 4213 |

**Table S3.** Colony counts for all four replicates of the inducible conjugation in the *trans* system. All plates were counted at the lowest dilution (pTAMob 2.0 + EP: $10^{-5}$, ΔTrbF + CP & ΔTrbC + CP: $10^{-3}$) plated apart from numbers with a star which were counted at a dilution of $10^{-1}$.

| | | Induced | Uninduced | Repressed |
|---|---|---|---|---|
| Replicate #1 | pTAMob 2.0 + EP | 39 | 50 | 52 |
| | ΔTrbF + CP | 174 | 0 | 0 |
| | ΔTrbC + CP | 105 | 0 | 0 |
| Replicate #2 | pTAMob 2.0 + EP | 106 | 90 | 25 |
| | ΔTrbF + CP | 204 | 3* | 1* |
| | ΔTrbC + CP | 145 | 3* | 9* |
| Replicate #3 | pTAMob 2.0 + EP | 1 | 18 | 43 |
| | ΔTrbF + CP | 209 | 0 | 0 |
| | ΔTrbC + CP | 132 | 0 | 0 |
| Replicate #4 | pTAMob 2.0 + EP | 46 | 220 | 159 |
| | ΔTrbF + CP | 28 | 1* | 1* |
| | ΔTrbC + CP | 50 | 1* | 1* |

**Table S4.** Colony counts for all six replicates of the inducible conjugation in the *cis* system. All plates were counted at the lowest dilution (pSC5GGv1: $10^{-5}$, pSC6: $10^{-3}$) plated apart from numbers with a star which were counted at a dilution of $10^{-1}$.

| | | Induced | Uninduced | Repressed |
|---|---|---|---|---|
| Replicate #1 | pSC5GGv1 | 465 | 2855 | 1002 |
| | pSC6 | 448 | 2* | 0 |
| Replicate #2 | pSC5GGv1 | 2055 | 921 | 1148 |
| | pSC6 | 132 | 1* | 0 |
| Replicate #3 | pSC5GGv1 | 968 | 1210 | 1692 |
| | pSC6 | 178 | 76* | 0 |
| Replicate #4 | pSC5GGv1 | 639 | 534 | 1984 |
| | pSC6 | 175 | 0 | 0 |
| Replicate #5 | pSC5GGv1 | 144 | 123 | 41 |
| | pSC6 | 39 | 17* | 8* |
| Replicate #6 | pSC5GGv1 | 116 | 62 | 56 |
| | pSC6 | 16 | 2* | 12* |
